# Regrowing the growth zone: metamorphosis kickstarts regeneration in the annelid, *Capitella teleta*

**DOI:** 10.1101/2025.08.05.668690

**Authors:** Alicia A. Boyd, Elaine C. Seaver

**Affiliations:** Whitney Laboratory for Marine Bioscience, University of Florida, 9505 Ocean Shore Blvd., St. Augustine, FL 32080-8610

**Author notes:** Author contributions: Alicia A. Boyd and Elaine C. Seaver conceived and designed the experiments. Alicia A. Boyd performed the experiments. Alicia A. Boyd and Elaine C. Seaver analyzed the data. Elaine C. Seaver and Alicia A. Boyd contributed reagents, materials, and analysis tools. Alicia A. Boyd and Elaine C. Seaver wrote the paper.

**Keywords:** Regeneration, metamorphosis, developmental plasticity, *Capitella teleta*

## Abstract

The ability to regenerate can vary across an animal’s life history. We previously showed that *Capitella teleta*, an annelid worm, gains regenerative ability with age. Although larvae do not replace lost structures, juveniles and adults regenerate posteriorly following metamorphosis. To determine whether metamorphosis enables juveniles to regrow structures lost as larvae, we amputated *C. teleta* larvae, removing posterior segments, the hindgut, posterior growth zone, anus, and pygidium. Metamorphosis was then induced in these amputated larvae and reared as juveniles for 3-, 7-, or 14-days. New growth in juveniles was assessed by confocal microscopy, EdU staining, immunohistochemistry, and nuclear staining. A pgz and new segments were observed by 7 days post metamorphosis. Morphallaxis of the digestive system in preexisting tissue was also observed. In addition, re-amputation of juveniles resulted in regeneration of the pgz, segments, and hindgut. Our results demonstrate that amputated *C. teleta* larvae can metamorphose into functional juveniles capable of growing new segments, suggesting that metamorphosis acts as a switch to enable regeneration of structures essential for growth. This study highlights the impact of metamorphosis on changes in developmental plasticity.

**Summary statement:** The annelid *Capitella teleta* can regain structures lost as a larva only after metamorphosis into a juvenile, indicating that metamorphosis may function as a switch in regeneration potential.

## Introduction

Regeneration is the ability to replace lost tissues and structures. The capacity an animal has to regenerate can change, even drastically, over the course of its life. Classic examples include amphibians, ascidians, and echinoderms (Bothe and Fröbisch, 2024; Byrnes, 1904; Carnevali, 2006; Jeffery, 2015a; Jeffery, 2015b; Phipps et al., 2020). Even humans have a correlative relationship between regeneration potential and age; as infants, humans can regenerate tissues, including digits, but this ability is progressively lost with increasing age. Unlike humans, many of the systems in which this dynamic has been demonstrated have an indirect life cycle. Animals that undergo indirect development generate larval forms before metamorphosing into a final adult form (Vervoort, 2011). Metamorphosis is a pivotal developmental process marked by the presence of a functional, free-living larva that in many marine invertebrates, rapidly transitions into a juvenile that loses larval-specific structures but maintains the functionality of adult structures generated as larvae and retained through metamorphosis (Hadfield, 2000; Hadfield 2001).

As such, the larvae of indirect developers can be surveyed for their regeneration potential and compared to their regeneration abilities in adult stages. For example, in anuran amphibians, such as *Xenopus laevis*, larvae are capable of regenerating multiple tissues and structures, including limbs. This ability is lost in adults, after metamorphosis (Edwards-Faret et al., 2021; Gibbs et al., 2011; Harrison, 1898; Phipps et al., 2020; Slack et al., 2008). In contrast, the ascidian *Ciona intestinalis* cannot regenerate before metamorphosis (i.e., as embryos or larvae), but can regenerate afterwards as juveniles and young adults; however, this ability is progressively lost with advanced age (Jeffery, 2015a; Jeffery, 2015b). Other animals, such as some crinoids, display ‘limited regeneration’ as larvae, only to display robust regenerative ability following metamorphosis (Carnevali, 2006; Carnevali et al., 1993; Vickery et al., 2001). Thus, for some animals, there is a loss of regenerative ability after metamorphosis (e.g., amphibians), while for others there is a gain of regenerative ability (e.g., ascidians and crinoids). For many indirect developers, it appears that metamorphosis is an event separating differences in regenerative ability.

*Capitella teleta* is an annelid worm that also displays different regeneration potentials before and after metamorphosis. *C. teleta* has robust posterior regenerative ability as juveniles and adults but shows very limited regulation for cell loss as embryos (Amiel et al., 2013; Pernet et al., 2012). We recently characterized the regenerative potential of *C. teleta* larvae and demonstrated that they have limited regenerative potential compared to that in juveniles and adults (Boyd and Seaver, 2023). Features reminiscent of the initial stages of juvenile regeneration occur, namely wound healing, cellular proliferation, expression of stem cell marker genes at the wound site, nerve extension into the wound, and neural specification. However, neural differentiation or the replacement of structures was not observed. It appears that *C. teleta* larvae can initiate the regeneration process but cannot successfully complete it, suggesting that *C. teleta* gains regenerative ability with age (Fig. 1A).

**Figure 1.**
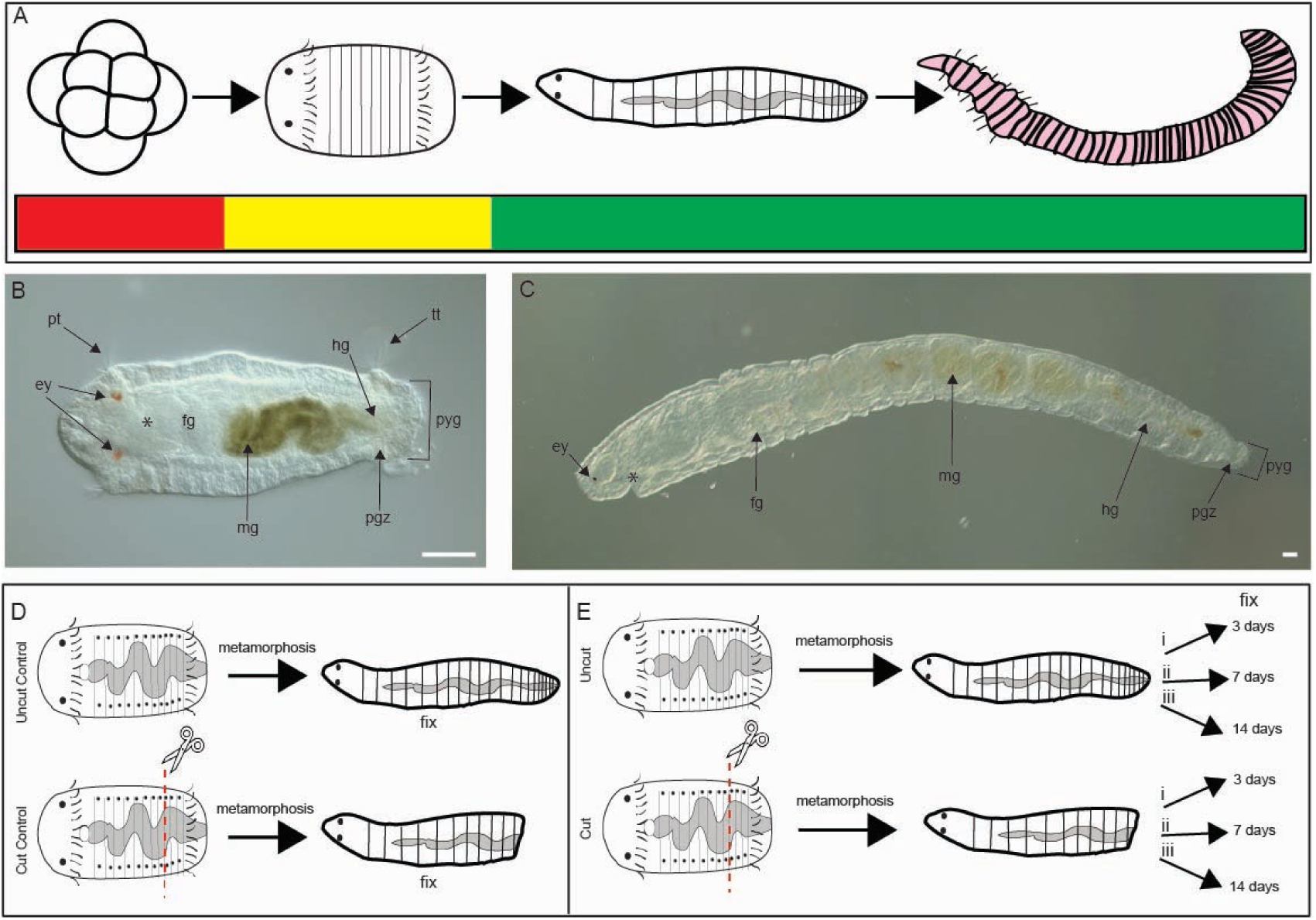
Evaluation of *Capitella teleta* regeneration following larval amputation and metamorphosis into juveniles. Anterior is to the left for all panels except embryo in (A). (A) *Capitella teleta* gains regeneration ability during development. Shown are schematics of representative developmental stages from left to right: embryo, larva, juvenile and adult. Red denotes lack of replacement of lost tissues (embryos), yellow denotes partial replacement (larvae) and green denotes stages that undergo complete regeneration (juveniles and adults). (B) Anatomical features of larva (Stage 9) and (C) two-week-old juvenile. Larva is in ventral view and juvenile is oriented in lateral view with ventral down. (D and E) Workflow of regeneration experiment. (D) Schematic showing manipulation of controls fixed and analyzed immediately following metamorphosis, either uncut or cut as larvae. (E) Schematic showing experimental design of juveniles cut, or uncut, as larvae, induced to undergo metamorphosis and raised for (i) 3 days, (ii) 7 days, or (iii) 14 days. (ey) eye, (pt) prototroch, (mg) midgut, (hg) hindgut, (pgz) posterior growth zone, (tt) telotroch, (pyg) pygidium. Amputation site denoted by scissors and red dotted line. Scale bars are 50 µm.

The presence of multiple morphological features of *C. teleta* can be leveraged to further study this dynamic phenomenon. The segmented body of *C. teleta* has complex structures and organ systems, including a central nervous system, muscles, and a regionalized digestive system. The ventral nerve cord is comprised of a single ganglion in each segment, and bilateral pairs of peripheral nerves exit each ganglion. Other segmentally repeated structures include the chaetae and segmental septae. Together, these structures enable the identification of individual segments and an ability to distinguish among different body regions. Furthermore, juvenile and adult animals of *C. teleta* demonstrate indeterminate growth by continually generating new segments (Seaver and De Jong, 2021). New segments are continually generated by the posterior growth zone (pgz), a subterminal region located immediately anterior to the terminal pygidium (Fig. 1B, C). The pgz is characterized by proliferating, undifferentiated cells and the expression of stem cell markers, such as *vasa* and *piwi* (Dill & Seaver, 2008; Seaver & de Jong, 2021). Larvae generate 13 segments before metamorphosis and have a pgz that express stem cell genes (Seaver, 2016). *C. teleta* juveniles and adults can fully regenerate each of these structures in the posterior direction within seven days and then continue the process of indeterminant growth (de Jong and Seaver, 2016). In contrast, amputated larvae cannot regenerate, although these larvae successfully metamorphose into juveniles following exposure to a metamorphic cue (Boyd and Seaver, 2023). Together, these features provide an opportunity to study regeneration on either side of the metamorphic divide.

Here, we ask whether amputated larvae can successfully replace their lost tissues following metamorphosis as juveniles, or whether regenerative ability is irrevocably lost. To address these questions, *C. teleta* larvae were amputated, induced to undergo metamorphosis, and evaluated for regenerative potential as juveniles. Over two weeks, we characterized cellular proliferation, generation of multiple tissues such as digestive system subregions as well as formation of new segments. In exploring this topic, our goal was to better understand the relationship between developmental maturation and regenerative potential. This study contributes to our understanding of the impact that metamorphosis can have on the shift of regenerative potential.

## Results

### New segments in amputated animals

We investigated whether animals amputated as larvae can grow new segments as juveniles. *C. teleta* larvae can be induced to undergo metamorphosis with either vitamin B or marine sediment (Burns et al., 2014). We previously demonstrated that amputated, competent larvae successfully metamorphose into juveniles when exposed to vitamin B (Boyd and Seaver, 2023). Following removal of the posterior 1/3 of the body by amputation and subsequent induction of metamorphosis, juveniles were scored for addition of segments at the following time points: immediately following metamorphosis (time zero’) (Fig. 1D), 3 days post metamorphosis (dpm), 7 dpm, and 14 dpm (Fig. 1E). Animals were scored for number of newly generated segments using three different markers, each scored independently. The three markers were the presence of ganglia in the ventral nerve cord (vnc) using a nuclear stain, paired peripheral nerves using anti-acetylated tubulin, and chaetae using both stains. The number of newly generated segments was compared between age matched cut and uncut animals at different time points after metamorphosis.

Over the course of fourteen days, we observed the appearance of new tissue and segment addition in both uncut and cut juvenile worms (Fig. 2A-Z). Juveniles fixed immediately after metamorphosis had approximately 13-14 segments in uncut control animals (time zero) (Fig. 2 A, C, E), whereas cut worms had approximately 7-10 segments and lacked a pygidium (Fig. 2 B, D, F). By 3 dpm, uncut juveniles had, on average, grown one new segment (Fig. 2G, I, K). In contrast, only an unsegmented mass of tissue distal to the amputation site was observed at 3 dpm in cut worms (Fig. 2H, J, L; n = 102/102). Multiple neurites continuous with the connectives of the ventral nerve cord were present in the new mass of tissue (Fig. 2L; n = 55/65). No discernable ganglia or peripheral nerves were observed in the new tissue (Fig. 2J, L). New segments, as indicated by presence of ganglia and peripheral nerves, were observed by 7 dpm in both cut and uncut worms (Fig. 2M-R). The width and length of these segments was visually smaller than those in preexisting tissue. The ventral nerve cord was present throughout the new tissue, with nerves spanning the pygidium (Fig. 2Q, R). The number of new segments in cut worms at 7 dpm ranged from 0 to 6, with an average of 4.5 new segments (n = 65). This was comparable to uncut worms, where the number of new segments ranged from 1 to 6, with an average of 4 new segments (n = 66). At 14 dpm, cut worms generated between 0 to 12 segments, with an average of 10.5 segments generated since metamorphosis (Fig. 2T, V, X, Z; n = 28); uncut worms grew an average of 10 new segments over 14 days (Fig. 2S, U, W, Y; n = 35). We also observed that the posterior-most tissue in cut worms resembled a reformed pygidium. In uncut juveniles, the pygidium is positioned distal to the pgz, resulting in a constriction of tissue width before flaring out into a bulb of tissue. The terminus of the pygidium frequently has an indentation marking the position of the anus (Fig. 2 S, U, W, Y). These features are present in juveniles amputated as larvae by 14 dpm (Fig. 2 T, V, X, Z). In summary, animals amputated as larvae and analyzed as 14-day old juveniles grow similar number of new segments to the number of segments generated in uncut worms.

**Figure 2.**
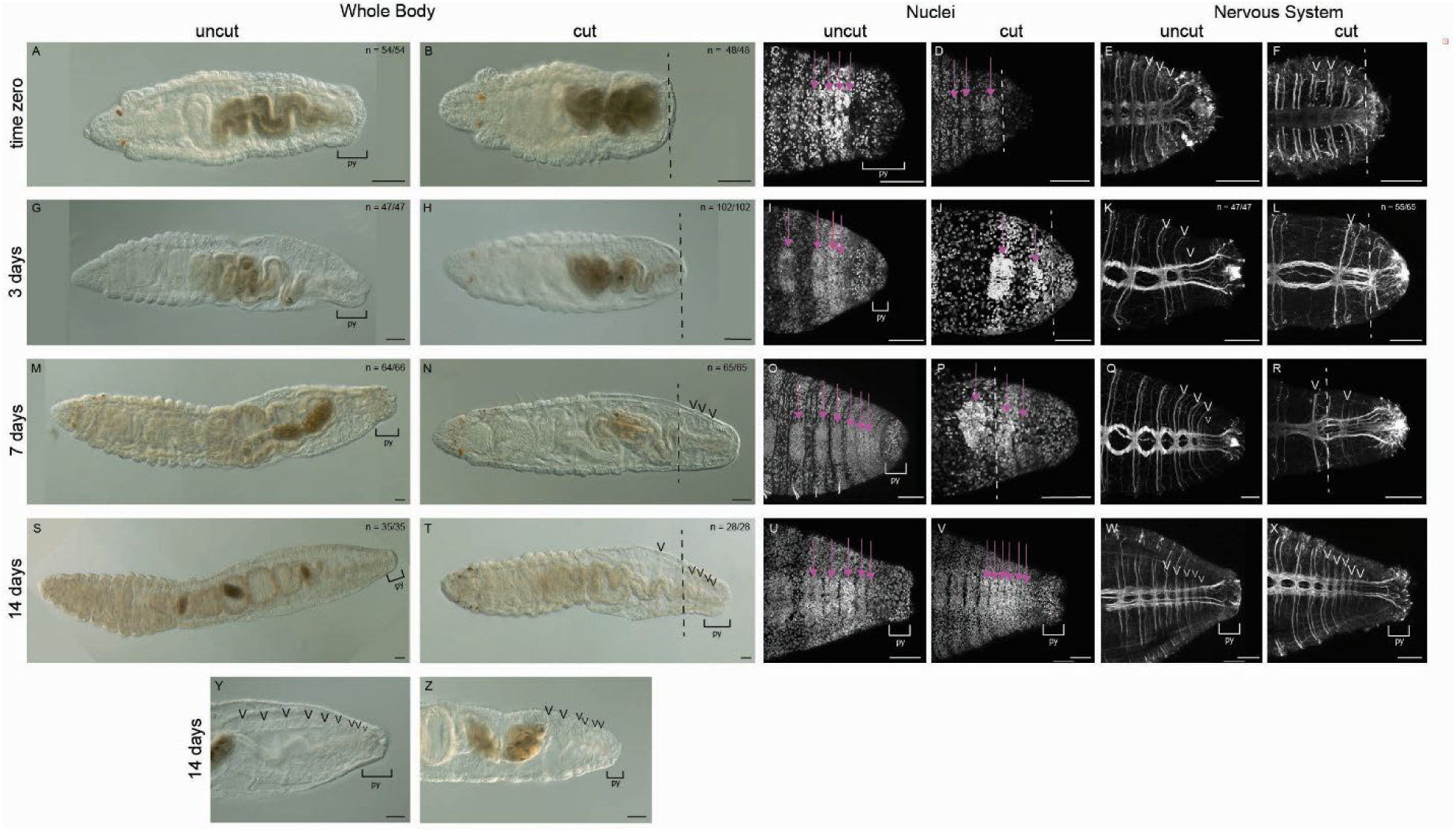
Juveniles amputated as larvae continually grow new segments. All images are oriented in ventral view, with anterior to the left. (A-F) Controls fixed immediately following metamorphosis (time zero). Juveniles fixed at 3 days post-metamorphosis (dpm) (G-L), 7 dpm (M-R) and 14 dpm (S-Z). (Y-Z) Magnified views of posterior ends of 14 dpm juveniles. (A-B, G-H, M-N, S-T, Y-Z) are transmitted light images. Arrowheads indicate segmental septae. (C-F, I-L, O-R, U-X) are stacks of confocal images. (D-E, I-J, O-P, U-V) Hoescht nuclear stain. Arrows indicate ganglia of the ventral nerve cord. (E-F, K-L, Q-R, W-X) Anti-acetylated tubulin antibody labeling of the nervous system. Arrowheads indicate pairs of peripheral nerves. Amputation site is represented by the dotted line. Black bracket indicates the pygidium. N denotes the number of animals scored that resemble the representative image. Arrowheads indicate individual segments by pointing to segmentally repeated structures. (py) pygidium. Scale bars are 50 µm.

### Regrowing the growth zone

The pgz is characterized by undifferentiated, proliferating cells and expression of stem cell markers in a band anterior to and immediately adjacent to the pygidium (Fig. 3A; n = 42/54) (Dill and Seaver, 2008; de Jong & Seaver, 2016). We examined expression of the stem cell marker *vasa* and EdU incorporation as indicators of a posterior growth zone in the newly formed tissue of cut and uncut worms. In amputated larvae, the pgz is removed, and EdU+ cells are not detectable at the wound site when animals are examined immediately after metamorphosis (Fig. 3B; n = 48/48). At 3 dpm, uncut worms have a distinct subterminal band of EdU+ cells, anterior to the pygidium (Fig. 3C; n = 46/47). In cut worms at 3 dpm, there are numerous EdU+ cells distal to the cut site that span the new tissue from the cut site to the posterior end of the animal and are present in the ectoderm and mesoderm (Fig. 3D; n = 95/102). By 7 dpm, a subterminal band of EdU+ cells is present in both uncut (Fig. 3E; n = 64/66) and cut worms (Fig. 3F; n = 63/63), indicating that the pgz has reformed in cut animals. To provide additional support to the assertion that this band of EdU+ cells is a pgz, we examined expression of the pgz marker *vasa* in 7 dpm worms (Fig. 4) (Dill and Seaver, 2008). As expected, the expression domain of *vasa* is missing from the posterior end of amputated animals immediately after metamorphosis, consistent with removal of the pgz during amputation (Fig. 4A; n = 14/14). In uncut juveniles, *vasa* is expressed in the ectoderm and mesoderm in a subterminal domain (Fig. 4B; n = 7/10). In 7 dpm juveniles that were amputated as larvae, there is novel subterminal *vasa* expression (Fig. 4C; n = 9/9), confirming the re-establishment of the pgz in juveniles.

**Figure 3.**
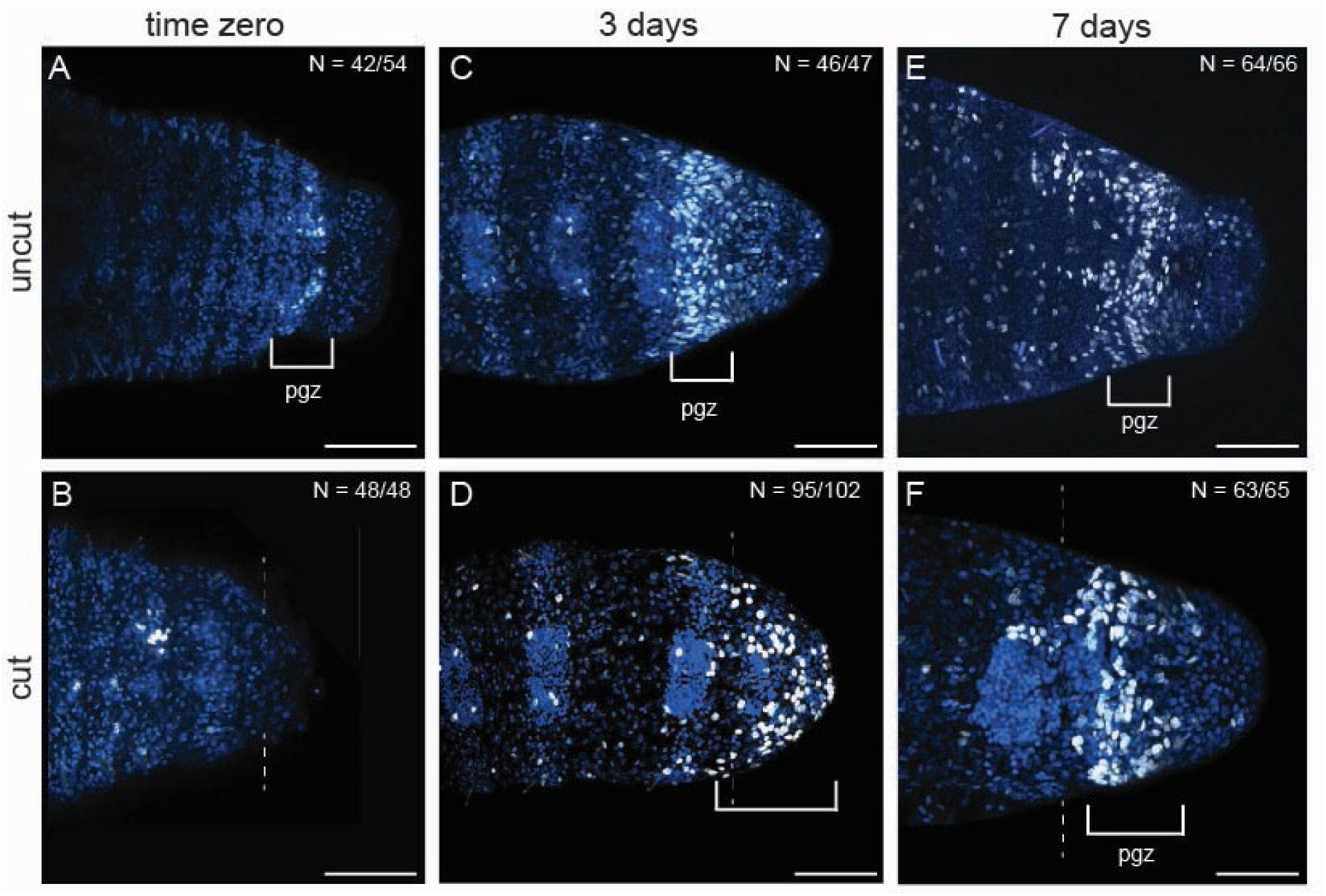
Posterior growth zone is reestablished in amputees by 7 days post metamorphosis. All images are oriented in ventral view, with anterior to the left. (A, C, E) are uncut juvenile controls and (B, D, F) are experimental juveniles that were cut as larvae, underwent metamorphosis and raised for various time intervals. (A) and (B) are animals fixed immediately following metamorphosis. (C) and (D) are 3 days post metamorphosis. (E) and (F) are 7 days post metamorphosis. White brackets indicate localized area of proliferating cells. Blue is Hoescht staining. White nuclei indicate EdU incorporation. Dotted line indicates the amputation site. N denotes the number of animals scored that are similar to the image represented. (pgz) posterior growth zone. Scale bars are 50 µm.

**Figure 4.**
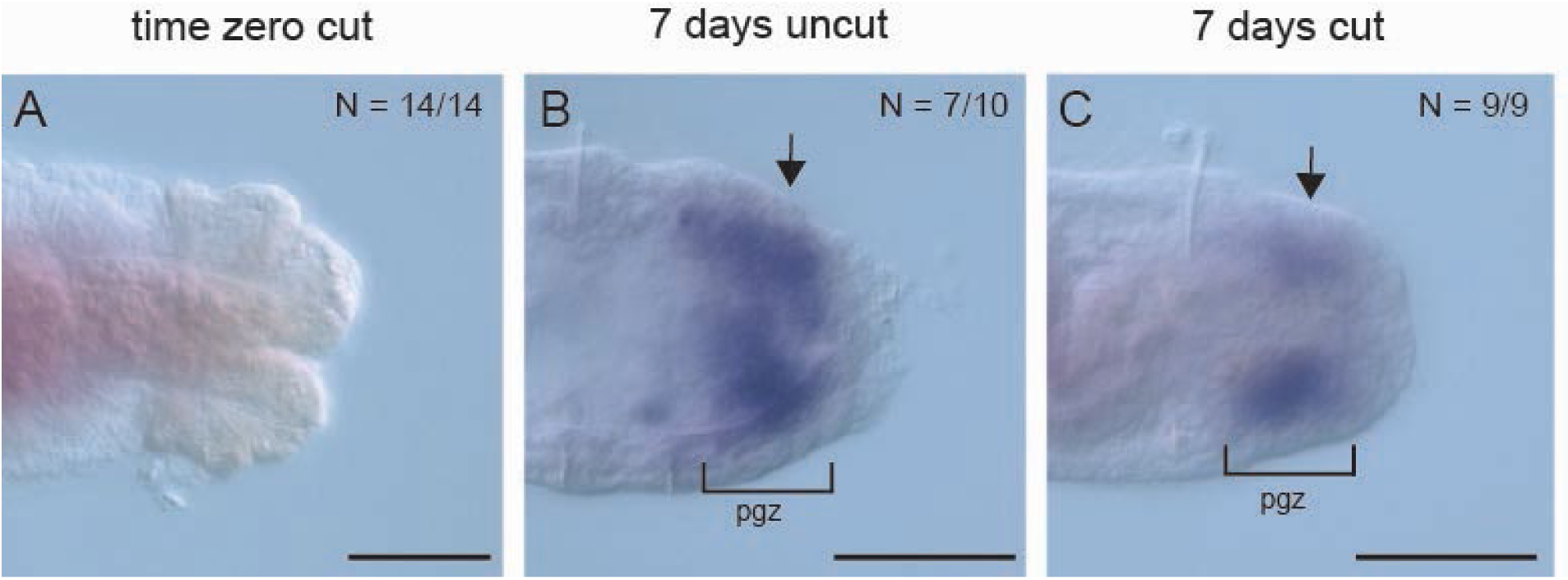
Posterior growth zone marker *vasa* is expressed in amputees by 7 days post metamorphosis. All images are of the posterior end of juveniles oriented in ventral view, with anterior to the left. (A) Cut time zero, immediately following amputation and metamorphosis. (B) Uncut 7-day old juvenile control, (C) 7-day old juvenile that was cut as a larva. Black brackets and arrows indicate subterminal domain of purple precipitate indicative of *vasa* expression by in situ hybridization. This expression pattern is typical of the posterior growth zone (compare B and C) (Dill and Seaver, 2008). N denotes the number of animals scored that are similar to the image represented. (pgz) posterior growth zone. Scale bars are 50 µm.

### Gut regeneration and morphallaxis in amputated animals

The digestive system of *C. teleta* is regionalized and broadly divided into a foregut, midgut, and hindgut. These regions can be identified by the presence or absence of ciliation in the lumen of the digestive tract, as visualized by anti-acetylated tubulin immunohistochemistry, and are clearly distinguishable by the time metamorphosis is complete. Cilia line the foregut and hindgut lumen but are absent from the lumen of the midgut (Fig. 5A, B; n = 47/47). Amputations in the midgut provides the advantage of being able to distinguish between midgut and hindgut identity in regenerating tissues.

**Figure 5.**
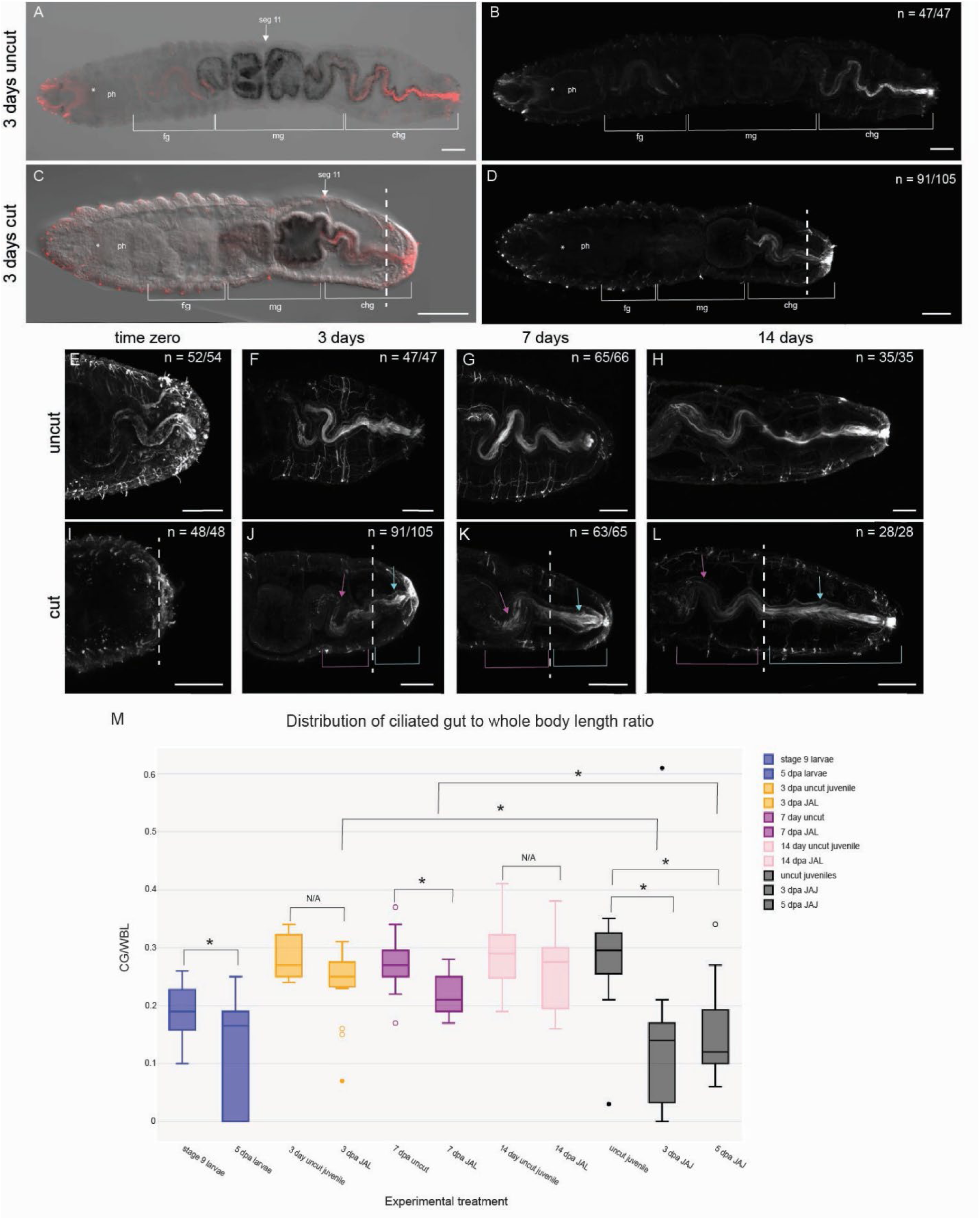
Posterior regeneration includes generation of ciliated hindgut in new tissue and repatterning of preexisting tissue. All images are oriented in ventral view, with anterior to the left. (A-B, E-H) are uncut juveniles, (C-D, I-L) are juveniles amputated as larvae. (A-B) are the same animal and (C-D) are the same animal. Brackets indicate subregions of the digestive system. The white arrows in (A and C) indicate the location of segment 11. Juveniles were fixed either immediately following metamorphosis (E, I) (time zero animals), 72 hours post metamorphosis (F, J), 7 days post metamorphosis (G, K) or 14 days post metamorphosis (H, L). Ciliation is marked with anti-acetylated tubulin antibody labeling and appears red in (A, C) and white in (B, D, E-L). Magenta arrows and brackets indicate ciliated hindgut in preexisting tissue. Teal arrows and brackets indicate ciliated hindgut in regenerated tissue. (M) Box and whisker plot showing proportion of the body length composed of ciliated hindgut for each experimental group. Y axis plots the ciliated hindgut: whole body ratio. CG, ciliated gut; WL, whole body length. X axis includes each experimental group. Uncut and cut animals for each experimental group are color coordinated: amputated larvae are blue, 3-day JAL are yellow, 7-day JAL are purple, 14-day JAL are pink, and JAJ, both 3 -and 5-day, are black. (JAL) juveniles amputated as larvae, (JAJ) juveniles amputated as juveniles. Brackets indicate pairings for statistical comparisons. (N/A) not statistically different. Asterisk indicates statistical significance. (A – D) Asterisk denotes the mouth. White dotted line indicates the approximate amputation site. N denotes the number of animals scored that are similar to the image represented. (ph) pharynx, (fg) foregut, (mg) midgut, (chg) ciliated hindgut. Scale bars are 50 µm.

From time zero through 14 dpm in uncut animals (Fig. 5E-H), a clear delineation can be made between the non-ciliated midgut and ciliated hindgut. Cut animals examined immediately after metamorphosis (time zero) are missing posterior ciliation compared to uncut animals of the same age (Fig. 5E, n = 52/54), as the hindgut was removed during amputation (Fig. 5I; n = 48/48). By 3 dpm, a new ciliated hindgut is observed in cut animals (Fig. 5J; n = 91/105), although it is abbreviated in length relative to the uncut controls (Fig. 5F; n = 47/47). The new ciliation in cut animals appears in pre-existing tissue proximal to the cut site as well as in the new tissue distal to the cut site (Fig. 5F, J). As illustrated in whole body images, the anterior boundary of ciliation in cut animals at 3 dpm is in a segment (e.g., segment 11, Fig. 5C, D) that in an uncut animal has midgut identity and therefore lacks cilia (Fig. 5A, B). This result indicates a remodeling of pre-existing tissue.

To better characterize the gradual replacement of hindgut following larval amputation, measurements of the lengths of the ciliated hindgut and the length of the body were made. From these measurements, the proportion of the body constituted by the ciliated hindgut was calculated and compared between cut and uncut animals of similar ages following metamorphosis (Fig. 5M). These proportions were also calculated in the context of successful posterior regeneration in juveniles amputated as juveniles (JAJ).

The proportion of the body that the ciliated hindgut constitutes shows a statistically significant difference between 14 dpm uncut control animals and juveniles cut as juveniles at both 3 dpa and 5 dpa (Fig. 5M and Table 1; Mann Whitney U test). This comparison was used as a framework for understanding how the hindgut regenerates in juveniles amputated as larvae, when compared with uncut juveniles of the same age. We hypothesized that the ratios would be different between uncut juveniles and juveniles amputated as larvae (JAL) at 3 dpm and 7 dpm, with the hindgut gradually regenerating and recovering normal proportional length with respect to the whole body. Surprisingly, we found that by 3 dpm, when ciliation first appears in the lumen of cut animals, the total proportion of the body comprised by ciliated hindgut is similar between uncut and cut animals and the difference is statistically insignificant (Table 1). This could be an artifact of linear measurements, as the gut in uncut animals has a more sinusoidal shape. As the uncut animals continue to grow with the addition of new segments, the ciliated hindgut elongates (Fig. 5G, H). This trend was also seen in cut animals (Fig. 5K, L). As expected, the difference in the ratio of the length of hindgut-to-whole-body between cut and uncut animals at 7 days is statistically significant (Fig. 5M, Table 1). This result indicates that the length of the hindgut is proportionally smaller in cut animals than in uncut animals (Fig. 5M). Over the course of two weeks, the hindgut of cut animals continues to elongate, and by 14 dpm, the proportion of the hindgut to whole-body length is similar between cut and uncut animals and is statistically insignificant (Fig. 5K, L, M, Table 1). The similarity observed at 14 dpm indicates a complete replacement of hindgut in cut animals resulting from a combination of morphallaxis of preexisting tissue and generation of hindgut in new segments.

**Table 1.** Mann Whitney U statistical analysis of hindgut length: whole body length ratio between various experimental groups. For each analysis, the z score and p value are listed.

| Experimental type | Groups compared | z score | p value | Significance |
| --- | --- | --- | --- | --- |
| Juveniles amputated as larvae (JAL) | 3 day uncut & 3 dpa cut | 1.7994 | 0.07195 | NOT significant |
|  | 7 day uncut & 7 dpa cut | 3.5959 | 0.0003233 | Significant |
|  | 14 day uncut & 14 dpa cut | 1.2441 | 0.2135 | NOT significant |
| Juveniles amputated as juveniles (JAJ) | 14 day uncut & 3 dpa cut | 3.2054 | 0.001349 | Significant |
|  | 14 day uncut & 5 dpa cut | 3.0567 | 0.002238 | Significant |
| Amputated larvae (AL) | St 9 uncut & 5 dpa larvae | 2.1481 | 0.0317 | Significant |
| Cross group comparison | JAJ 3 dpa & JAL 3 dpa | -3.2375 | 0.001206 | Significant |
|  | JAJ 5 dpa & JAL 7 dpa | -3.3618 | 0.0007744 | Significant |

To evaluate whether hindgut regeneration occurs similarly between juveniles amputated as juveniles and juveniles amputated as larvae, the ratios of hindgut to whole body length compared between these two groups. There is a significant difference between JAJ at 3 dpa and JAL at 3 dpm as well as between JAJ at 5 dpa and JAL at 7 dpm (Fig. 5M, Table 1; Mann Whitney U test). These results indicate that regenerating juveniles recover the proportionality of the hindgut differently, depending on the developmental stage the initial amputation occurs.

Since the hindgut continues to elongate in cut and uncut animals as the juveniles continually grow and add new segments, we looked for dividing cells in the hindgut epithelium. In uncut juveniles, EdU+ cells are present in the epithelial lining of the hindgut at 3 dpm (Fig. 6A, A’). By 14 days, the number of EdU+ cells visually decreases in the hindgut epithelium (Fig. 6B, B’). In contrast, other surrounding tissues (e.g., pgz) do not show the same apparent decrease in EdU incorporation. In cut animals at 3 dpm, EdU+ cells are also present in the hindgut epithelium (Fig. 6 C, C’). However, EdU+ cells persist throughout the hindgut for two weeks (Fig. 6D, D’); this result contrasts with uncut animals of the same age. Across all samples, an aggregation of EdU+ nuclei occurs in the gut epithelium of the segment at the midgut/hindgut transition. The presence of dividing cells in the growing hindgut epithelium is consistent with a local cellular origin of new tissue.

**Figure 6.**
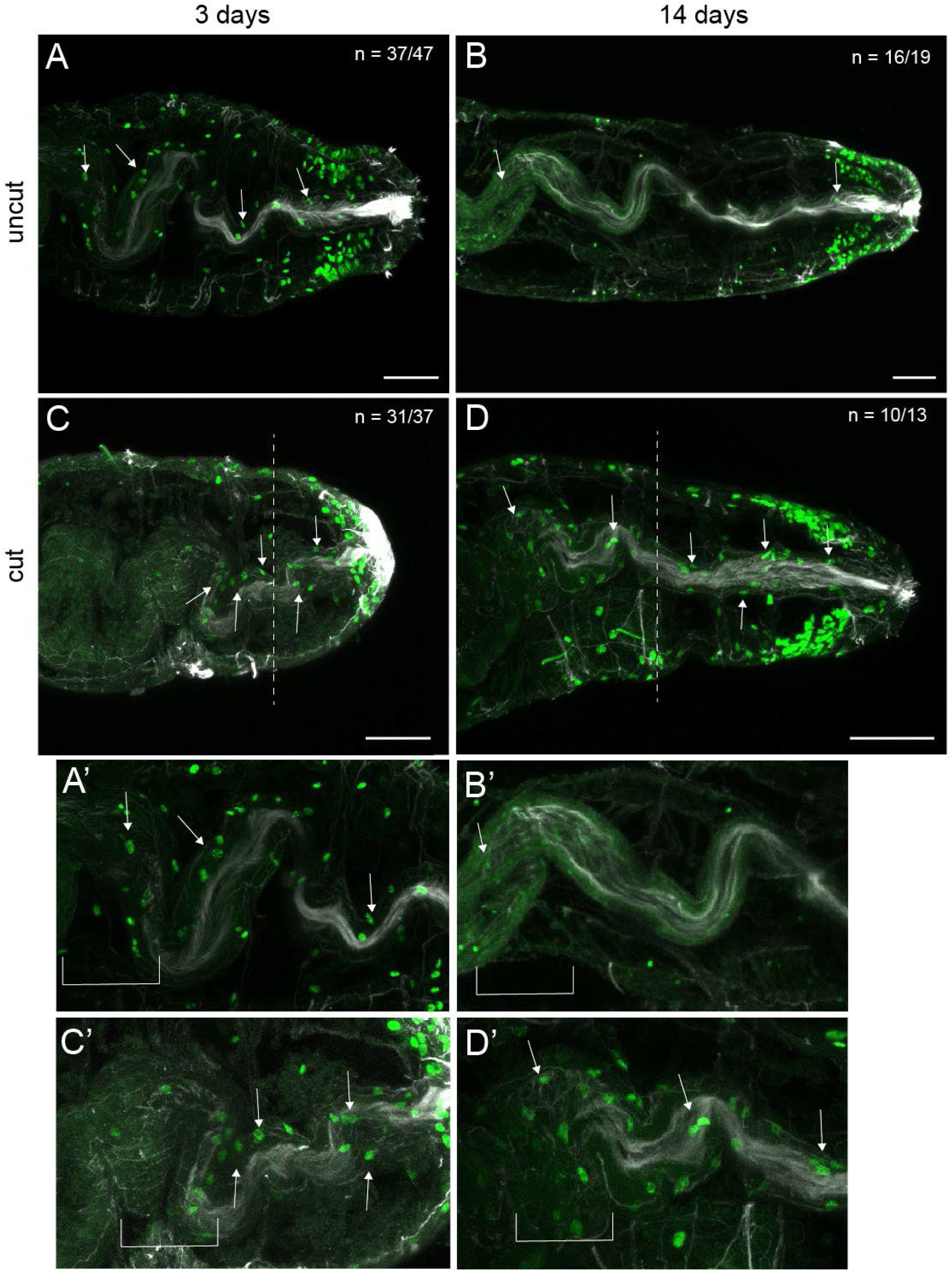
Elevated proliferating cells in the regenerating hindgut epithelium. All images are oriented in ventral view, with anterior to the left. (A-B) are uncut animals, (C–D) are juveniles cut as larvae. (A) and (C) are 3 days post-metamorphosis juveniles. (B) and (D) are 14 days post-metamorphosis juveniles. (A’-D’) are magnified images of (A-D), respectively. White arrows indicate EdU positive cells in the hindgut. EdU-positive nuclei are green. Ciliated hindgut is visible with anti-acetylated tubulin antibody labeling (white). White dotted lines denote approximate amputation site. White bracket indicates the region in which the midgut transitions to the hindgut. N denotes the number of animals scored that resemble the representative image. Scale bars are 50 µm.

To better understand whether morphallaxis of the hindgut is unique to juveniles, we also analyzed amputated larvae for reappearance of ciliation. In stage 6 larvae, the digestive system is still maturing and therefore does not have ciliation in the posterior portion of the digestive tract (Fig. 7A). The area in which the hindgut normally develops is removed during amputation (Fig. 7B). In late-stage larvae (stage 9, uncut), the ciliated hindgut is now present in the posterior end of the trunk and pygidium (Fig. 7C, C’; n = 13/13). By 5 dpa, half of the animals have gut ciliation, a statistically significant difference in comparison with uncut stage 9 larvae (Fig. 6M, Table 1; Fig. 7D, D’; n = 9/16). This ciliated region is shorter than what is observed in uncut age matched larvae and is present in the pre-existing tissue (Fig. 7D, D’). Therefore, despite removal by amputation of hindgut precursors, these observations show that morphallaxis occurs in amputated larvae, albeit more slowly and to a smaller degree than in amputated juveniles.

**Figure 7.**
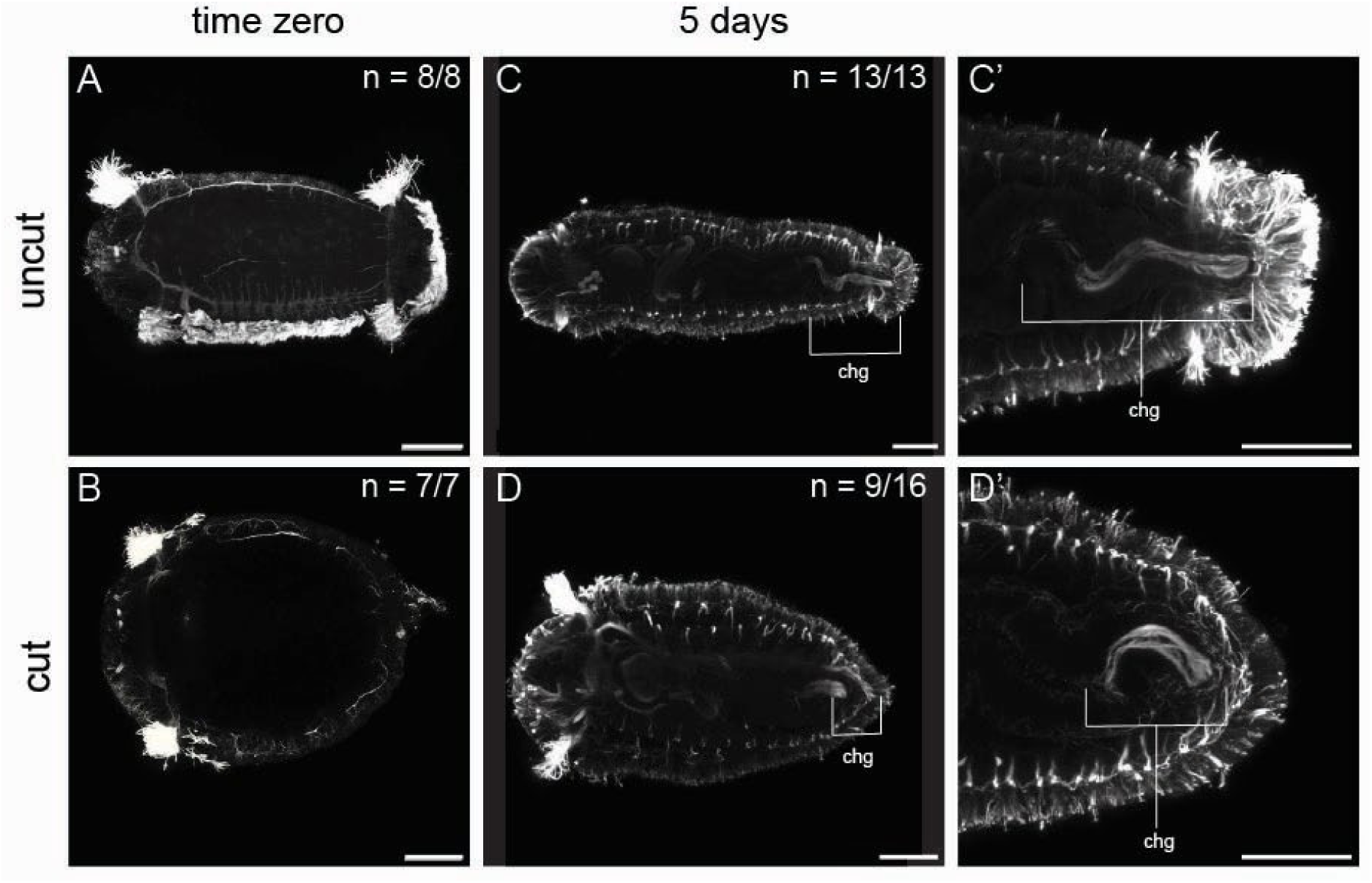
Amputated larvae develop ciliated hindgut. All images are oriented in ventral view, with anterior to the left. (A, B) Larvae fixed at stage 6 either following amputation (B) or as uncut controls (A) (time zero). (B) Cut animals were allowed to wound heal prior to fixation. (C, C’) Late-stage uncut larvae (Stage 9). (D, D’) Larvae 5 days post amputation. (C’, D’) are enlarged views of C and D, respectively. All images are larvae are labeled with anti-acetylated tubulin antibody (white). Brackets indicate the ciliated hindgut. N denotes the number of animals scored that resemble the representative image. (chg) ciliated hindgut. Scale bars are 50 µm.

### Re-amputation experiment

To evaluate if juveniles cut as larvae respond to amputation the same as previously unmanipulated juveniles, we amputated two groups of 2-week-old juveniles at segment 10 (Fig. 8A). One group was previously amputated as larvae and then again as juveniles, while the other group was only amputated one time as juveniles. Worms that were amputated as larvae and again as juveniles are referred to as ‘double cut’ and the worms cut only as juveniles are referred to as ‘single cut’ controls. The animals were scored for formation of new segments at 7 dpa. New segments were identified by presence and number of segments using the same characters as in the previous experiment, namely, newly formed ganglia and pairs of peripheral nerves distal to the cut site. 7 dpa is sufficient time for regeneration of a pgz and multiple new segments in 2-week-old juveniles, thus both features could be scored.

**Figure 8.**
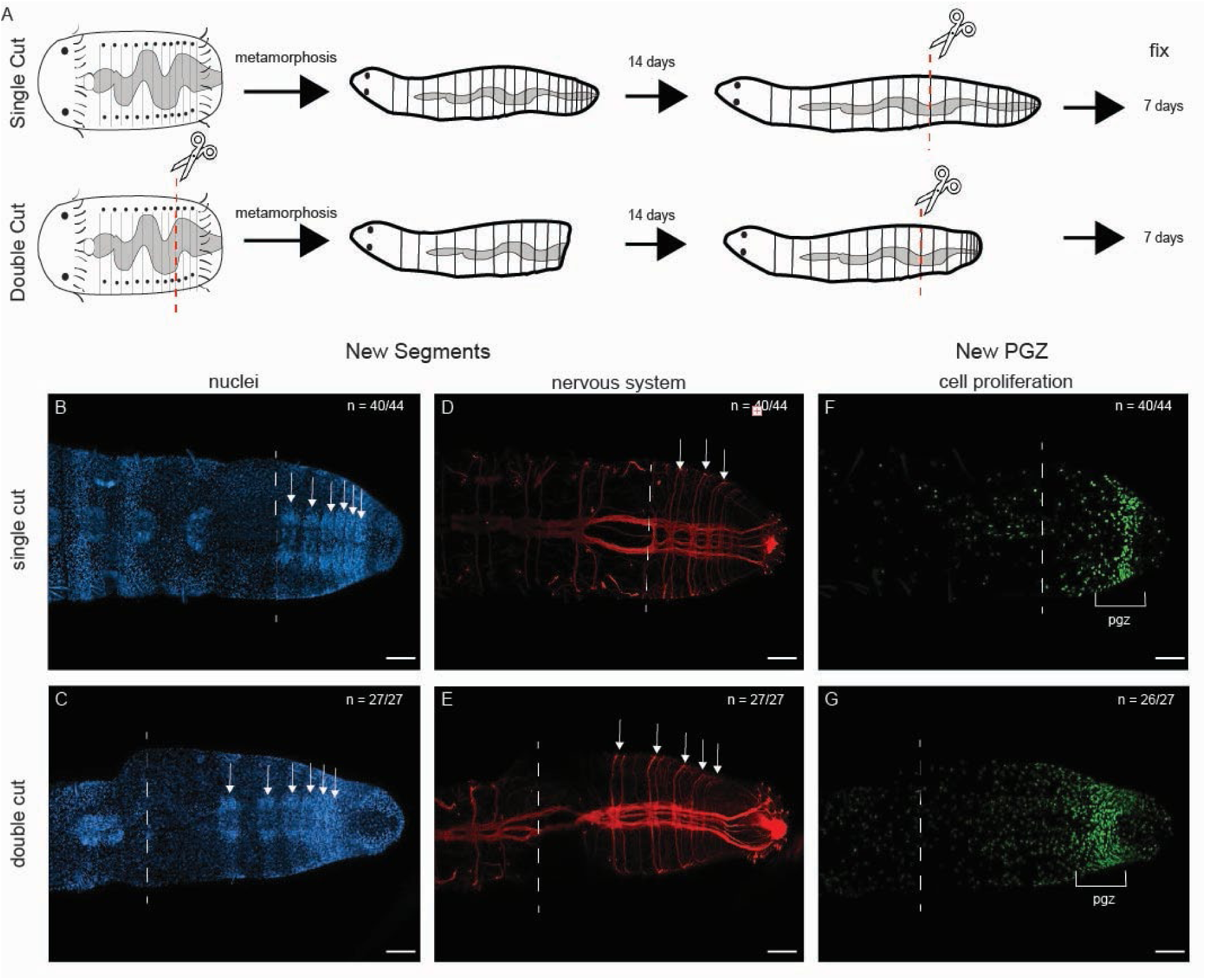
Successful regeneration following sequential amputations. All images are oriented in ventral view, with anterior to the left. (A) Schematics showing experimental design of the re-amputation experiment. Single cut denotes controls amputated only as juveniles. Double cuts denote animals amputated as larvae and again as juveniles. (B-G) Juveniles 7 dpm. (B-C) Nuclear staining highlights the segmental ganglia (arrows). (D-E) Anti-acetylated tubulin labeling shows paired peripheral nerves and ganglia of individual segments. (F-G) EdU incorporation showing the position of the pgz. Blue is nuclear staining; red is anti-acetylated tubulin antibody labeling and green nuclei indicate EdU incorporation. White arrows denote individual segments. The dotted line indicates the amputation site. The bracket marks the boundaries of the pgz. (pgz) posterior growth zone. Scale bars are 50 µm.

Both single cut and double cut worms regenerated after 7 days. Single cut worms grew up to 10 new segments; the average total length of the worms resulted in 17 total segments (Fig. 8B, D; n = 44). Double cut worms grew up to 12 new segments; the average total length of the worms resulted in 16 total segments (Fig. 8C, E; n = 27). New posterior growth zones were present in both uncut and cut worms as visualized by the presence of a subterminal band of EdU+ nuclei in single (Fig. 8F; n = 40/44) and double cut juveniles (Fig. 8G; n = 26/27). Ciliated hindguts were present in both single cut (n = 41/44) and double cut (n = 27/27) worms. Taken together, double cut worms regenerated to an extent comparable to the single cut worms.

## Discussion

### *Capitella teleta* replaces lost tissues only after metamorphosis

In this study, we demonstrate a gain in regeneration ability in the annelid *C. teleta* following the induction of metamorphosis into juvenile stages. We asked whether regenerative ability was inherent to specific life stages or could be influenced, even hindered, by a wounding event and tissue removal during a previous life stage. *C. teleta* were amputated during a regenerative incompetent stage, i.e. larva, and evaluated for the replacement of lost structures during a regenerative competent stage, i.e. juvenile. We found that juveniles derived from amputated larvae replaced the posterior growth zone, nerves and digestive system, and generate new, fully patterned segments following metamorphic induction. Juveniles added similar numbers of segments within the same time interval during regeneration and normal growth as uncut controls. These new tissues appear to be functional; the presence of fecal pellets in the digestive tract is consistent with the restoration of a functional digestive system (A.B., personal observation), and the continued addition of new segments is indicative of a functional pgz. Moreover, when amputated a second time as juveniles, the worms once again underwent successful regeneration. With these observations, it appears that amputation during larval stages does not hinder the regeneration ability of juveniles.

The ability to fully regenerate in *C. teleta* appears to be triggered by metamorphosis. In amputated larvae (AL), we only observed the beginning stages of posterior regeneration, including wound healing, cell proliferation and stem cell marker expression at the wound site and neural extensions into the wound, but not complete replacement of lost larval or adult structures such as ciliary bands, pgz or pygidium (Boyd and Seaver, 2023). In the vast majority of cases, we did not observe the replacement of lost structures, i.e. ciliary bands, pgz, pygidium, etc. However, as seen in this study, these animals can successfully replace the pygidium and pgz at a later stage following metamorphosis as juveniles. This process is initiated early, within three days of metamorphosis, indicating that cut animals are able to both recognize the absence of lost structures and initiate regeneration to replace these tissues in the juvenile body. It is noteworthy that, although extremely rare (i.e. in less than 10% of larvae), a few animals did regenerate the telotroch and pygidium. Similarly, in another study, in response to targeted deletion of the germline precursor in *C. teleta* embryos, all 2-week juveniles regenerate germline cells, while only 13% of larvae exhibit some degree of regeneration of germline precursors (Dannenberg and Seaver, 2018). Taken together, these data suggest that only a small fraction of larvae have the competency to receive and respond to the initial regeneration signal following amputation (i.e. wound heal, initiate cell proliferation, extend neurites, etc.), but cannot complete the regenerative process. We hypothesize the change in regenerative competency may be due to a lack of nutritional resources, an absence of a molecular signal, or an ability to respond to signals that are present post metamorphosis. We hypothesize that the change in regenerative competency is the result of both nutritional and molecular changes.

Our experimental manipulations temporally uncouple the events of regeneration by inducing metamorphosis directly after the amputation stimulus, thereby temporally separating surgical amputation from the completion of regeneration through the process of metamorphosis. *C. teleta* can thus successfully regenerate missing structures that normally persist through metamorphosis into juvenile stages and persist as adult structures. These observations suggest that the molecular and physiological signals initiated by amputation persist through the process of metamorphosis. Our results also demonstrate a temporal persistence between the initial wounding event and the delay in regeneration response following the metamorphic cue.

Our observations in *C. teleta* represent just one example of the varied regenerative potential observed in annelids. In posterior regeneration of many other annelids, the pgz and pygidium are the first structures to regenerate (Boilly et al., 2024). Our results in *C. teleta* are similar; we observe that the pgz and pygidium are formed in juveniles amputated as juveniles (JAJ) animals 4 dpa and in juveniles amputated as larvae (JAL) animals between 3 - 7 dpa. In both groups, the pgz appears to be functional, as demonstrated by the birth of new, patterned segments. The results of our re-amputation experiment in juveniles support previous observations that *C. teleta* adults are resilient to repeated amputation (Hill et al., 1986). The number of new segments generated by juveniles that were amputated as larvae is similar to uncut animals at 1- and 2-weeks post-amputation, suggesting that metamorphosis resets the regeneration program, independent of the number of amputation events. We also observed that the number of new segments generated was similar regardless of whether they were amputated once as larvae or amputated twice. In contrast, other annelids such as *Syllisamica quatrefages* and *Platynereis dumerilii* regenerate new segments at a significantly faster rate immediately following amputation (i.e. first 10 days) than in the proceeding weeks (Boilly et al., 2024; Metzger and Özpolat, 2024). In *S. quatrefrages*, the end of the initial regenerative response is marked by a return in growth rate to that of unamputated worms (Boilly et al., 2024). In summary, there are both similarities and differences in the specifics of posterior regeneration between our observations in *C. teleta* and those from other adult annelids.

Despite numerous surveys in annelids of juvenile and adult regeneration ability (Bely, 2006; Metzger and Özpolat, 2024; Starunov et al., 2020; Zattara and Bely, 2016), to our knowledge, *C. teleta* is the first annelid whose regeneration ability has been systematically characterized across the metamorphic transition. As annelids display a great variation in adult regenerative ability, it can be inferred that this variation extends to earlier developmental stages. Many annelids have larval stages that can be surveyed for regenerative potential. Characterization of additional stages can be conducted on species whose regeneration abilities have already been described as adults and be included in future surveys to gain a better understanding of how regeneration varies with maturation in annelids.

### Morphallaxis response in digestive tissue

We observed hindgut regeneration distal to the cut site in both amputated larvae (AL) and juveniles amputated as larvae (JAL) animals. Notably, we also observed new hindgut tissue proximal to the cut site within 3 dpa in regions that were previously midgut, indicating morphallactic remodeling of the old tissue, prior to new segment formation distal to the amputation site. We interpret this rapid tissue specific repatterning as being under evolutionary pressure to quickly replace all components of the digestive system in preparation for critical nutritional input. Furthermore, the hindgut continues to grow throughout adult life, ultimately recovering the same proportionality of hindgut to body length by 14 dpm. These data, in combination with previous observations of a limited shift in *Hox* gene expression domains following amputation (de Jong & Seaver, 2016), show that epimorphic and morphallactic processes occur concurrently in *C. teleta* regeneration. Similar remodeling has been observed in other annelids (Bely, 2014; Takeo et al., 2008; Zattara, 2020). For example, regionalization of the digestive system in *Enchytraeus japonesis*, as evidenced using histochemistry of alkaline phosphatase and gut specific markers, reestablishes proportionally following amputation (Myohara et al., 1999; Takeo et al., 2008). In annelids, pre-existing digestive and neural tissues can be remodeled within hours to days after amputation (Bely, 2014; Özpolat and Bely, 2016; Takeo et al., 2008).

Although gut remodeling occurs in *C. teleta*, the cellular source of the new hindgut, either proximal or distal to the cut site, is currently unknown. Our observations of EdU+ incorporation in the lining of the digestive tract in *C. teleta* might be indicative of a local cellular source (Fig. 6). Alternatively, hindgut in tissues proximal to the cut site may have a different cellular source from the regenerating hindgut distal to the amputation site. In other annelids such as *Lumbriculus variegatus*, cells in the gut endoderm may contribute to the regenerated gut (Tweeten and Reiner, 2012). Recent work in *Platynereis dumerilii* reports the presence of gut progenitor cells that contribute to regenerating tissues (Bideau et al., 2024). Outside of annelids, a study in the planarian *Schmidtea mediterranea* indicates that gut regeneration is the combined result of contribution from neoblast stem cells and remodeling of preexisting tissue (Forsthoefel et al., 2011). Future work is needed to explore whether lineage restricted stem cell precursors reside in the gut epithelium, or other cellular sources contribute to the regeneration of the digestive system in *C. teleta*.

### Metamorphosis as a key transition event in regenerative potential

Metamorphosis is an event marked by substantial morphological, physiological, and ecological changes to undertake a new body form (Bishop et al., 2006; Monaghan et al., 2014). Gain or loss of regenerative potential across the metamorphic divide appears to be one of the results of these substantive changes, including altered cell plasticity, molecular signaling, metabolic differences, or stem cell contribution (Booth et al., 2025). Molecular changes that occur during and immediately following metamorphosis include gene transcription, protein synthesis, chromatin accessibility, and hormone fluctuations. Metamorphosis can occur when transcription is repressed pharmacologically in 96% of *Phestilla sibogae* larvae, but new gene and protein activity are required in the novel juvenile structures (Hadfield, 2000). The timing between the initiation of metamorphosis and novel transcriptional activity is mostly unknown, although it is likely influenced by factors such as an increase in availability of nutrients, an acclimation to the new ecological setting, or epigenetic changes (Hadfield, 2000). Recent ATACseq studies have demonstrated differentially accessible regions of chromatin between pre- and post-metamorphic stages in flatfish and in the hemichordate *Schizocardium californicum* (Bump, 2022; Guerrero-Peña et al., 2023). In anurans, there is evidence that metamorphosis inhibits regeneration associated gene transcription, either directly or epigenetically (Christen and Slack, 1997; Monaghan et al., 2014; Phipps et al., 2020). Axolotls that are experimentally induced to undergo metamorphosis experience a reduction in regeneration rate and success (Monaghan et al., 2014). These results were interpreted to be due to cells taking longer to complete the cell cycle, with reduced cell proliferation. Hormonal changes can also influence regeneration outcomes. For example, an increase in thyroid hormone directly leads to a loss of regenerative potential in the central nervous system of *Xenopus laevis* (Gibbs et al., 2011). Taken together, our results that position metamorphosis as a key event in the shift of regeneration potential in *C. teleta* is shared by a range of metazoans.

The molecular, cellular, and structural changes associated with metamorphosis serve as a switch in regenerative potential. In some animals such as anurans, this switch hinders regeneration. However, in some indirect developing invertebrates (e.g. the crinoid *Antedon bifida* (Vickery et al., 2001), *C. teleta, etc.*), metamorphosis promotes the onset of regenerative success. The relationship between metamorphosis and regeneration may be, in part, due to similarities in their genetic components. Additionally, metamorphosis may result in tissue maturation that renders tissues receptive to signals required for regeneration. Metamorphosis and regeneration are both post-embryonic developmental processes that result in tissue reorganization, cell birth, apoptosis, and morphogenesis (Alibardi, 2024; Bothe & Fröbisch, 2023). It has been speculated that regeneration reuses many of the same genetic features of metamorphosis, which were lost in vertebrate adult organ and neural regeneration (Alibardi, 2019). While the process of metamorphosis across metazoans is highly diverse, there could nonetheless be an evolutionarily conserved pressure for a regeneration-associated switch, yet to be identified.

### Future Directions

Our hypothesis that a combination of changes in nutritional status and molecular inputs following metamorphosis resulting in increased regeneration ability can be analyzed by testing nutritional status and molecular signaling before and after metamorphosis. In regard to nutrition, it has been observed that *C. teleta* juveniles begin eating within 1-2 hours of metamorphosis. As the larvae are nonfeeding, this immediate nutritional input could provide the necessary energy to fuel completion of the remaining regeneration stages. It has also been shown that the extent of regeneration in juveniles depends on nutritional supply. It would be interesting to observe whether starving a newly metamorphosed, amputated juvenile 3 days post metamorphosis would affect the initial response observed in this study. If we observe the localized cell proliferation and/or re-establishment of a pgz, this could indicate a stage specific molecular cause rather than a requirement for additional nutritional input for these aspects of regeneration.

The observation that *C. teleta* larvae display early but not late stages of regeneration suggests that a signal critical for completion of regeneration might be absent. To that end, treating amputated larvae with an exogenous signal could yield an increase in regeneration progression and the formation of ciliary bands or a pygidium. Wnt signaling is important for regeneration in many animals, including planarians and annelids. For example, in the direct developing planarian *Schmidtea polychroa*, there is a gradual developmental increase in head regeneration (Booth et al., 2025). By altering canonical Wnt signaling via β-catenin-1 RNAi knockdown, head regeneration was induced precociously in 97% of animals, at a stage normally incapable of regeneration. Similarly, exposing *C. teleta* larvae to a Wnt signaling activator may promote completion of regeneration. Such experiments would provide support for the role of Wnt signaling in the regeneration program of *C. teleta*; this signal may not naturally be present at sufficient levels in larvae. If there is no change in regeneration potential following exposure to a Wnt activator, the receiving tissue may be immature and unreceptive to signaling or perhaps a different signaling pathway is required. Consistent with the idea that activation of canonical Wnt signaling might function at later stages of regeneration in *C. teleta*, a previous study in juveniles indicates that increasing cWnt signaling is not required to activate the earliest stages of regeneration (Kunselman and Seaver, 2025)

## Conclusions

Metamorphosis is a key event in the subsequent acquisition of successful regeneration in *C. teleta*. Our results indicate that posterior regeneration is inherent in post-larval stages following the appropriate metamorphic cue. Comparisons of regenerative potential in the same animal (e.g. with ontogenetic or physiological boundaries), can provide insights into the impediments and catalysts of regeneration (Goss, 1992). Additional sampling of regenerative ability at different life history stages in other species will provide insight into the onset of regeneration and its relationship with developmental maturation. Varied regeneration abilities across annelids allow us to better understand and parse out the degrees of plasticity that exist across the life cycle.

### Limitations of this study

Due to low recovery of re-amputation animals, only two independent experimental replicates were conducted for the re-amputation experiments. Measurements of the ciliated hindgut were done using linear measurements, although the gut is curvy. A better representation of the proportion of the hindgut to the worm’s entire length would be measuring the length of the hindgut’s curves. Limited molecular markers were used to characterize regenerating tissues.

## Methods and Materials

### Animal care

A colony of *Capitella teleta* was maintained in glass finger bowls with filtered sea water (FSW) and marine mud at 19°C, as previously described (Grassle & Grassle, 1976). Larvae were dissected from brood tubes of healthy females and raised in FSW until the appropriate age for amputation. Larvae from a single brood tube were divided between experimental, cut groups, and uncut control groups.

### Amputations

Larvae were transferred to a methyl cellulose solution (1.6% methyl cellulose: filtered sea water) on a 60 mm petri dish lid. Once immobilized, larvae were amputated using pulled glass pipettes (Boyd and Seaver, 2023), removing the posterior end of the larva. Stage 9 larvae were amputated for the regeneration experiments and stage 6 larvae for experiments on 3 dpa and 5 dpa larvae. Developmental stage was determined using a standard staging system (Seaver et al., 2005). The larvae for time zero (fixed following amputation), were transferred to filtered sea water for 5 hours to allow for wound healing to occur and then fixed in 4% paraformaldehyde for 1 hour at room temperature (EdU and antibody treatments) or overnight at 4°C (in situ hybridization).

Stage 9 larvae were amputated and allowed to wound heal for 6 hours. The amputated larvae induced to undergo metamorphosis (Boyd and Seaver, 2023) by either exposure to vitamin B in FSW (time zero animals) or transferred to a finger bowl with mud and FSW. Each group of time zero metamorphosed larvae were visually observed for successful metamorphosis, i.e. burrowing behavior, dropping of ciliary bands, and body elongation before fixation. Juvenile worms were fed with marine sediment and underwent water changes weekly until they were removed from the mud by sifting. The worms were placed in 0.5% cornmeal agar (Sigma Aldrich): FSW plates for 1–4 hours to remove attached debris. Thereafter, they were transferred to a petri dish with 1:1 MgCl: FSW solution for 15 minutes. In preparation for amputation, individual animals were moved to MgCl: FSW droplets on a platform of black dissecting wax (American Educational Products, Fort Collins, CO, USA). Worms were amputated using a microsurgery scalpel (Feather; 15-degree blade, Carlsbad, CA, USA). Using the presence of chaetae and segment indentation to count individual segments, worms were amputated at the boundary between segment 10 and 11. Following amputation, worms were placed in 1:1 MgCl: FSW solution for a 1–2 hour recovery. Worms were then placed in fresh bowls of FSW with a teaspoon of sieved estuarine mud for one week. After 1 week, individuals were sifted from the mud, placed into agar plates for debris removal, and then processed for either EdU incorporation, immunohistochemistry, or whole mount in situ hybridization.

For the re-amputation experiment, the initial amputation was conducted as previously described in this section. At 14 dpm the cut and uncut juveniles were sifted from the mud and placed in cornmeal agar plates for 1-4 hours to remove attached debris. In preparation for amputation, individual animals from either the cut or uncut groups were moved to MgCl: FSW droplets on a platform of black dissecting wax (American Educational Products, Fort Collins, CO, USA). Worms were amputated using a microsurgery scalpel (Feather; 15-degree blade, Carlsbad, CA, USA). Using the presence of chaetae and segment indentation to count individual segments, worms were amputated at the boundary between segment 10 and 11. Following amputation, worms were placed in 1:1 MgCl: FSW solution for a 1–2 hour recovery. After 1 week, individuals were sifted from the mud, placed into agar plates for debris removal, and then processed for either EdU incorporation or immunohistochemistry.

### Induction of metamorphosis

After amputation, animals were placed in glass finger bowls with a thin layer of marine mud and filtered seawater to induce metamorphosis. The animals were reared at 20°C until the desired time post metamorphosis. Animals were then removed from the mud and allowed to burrow in 35 mm plastic dishes half filled with 0.5% cornmeal agar (Sigma Aldrich): FSW for 1–4 hours to remove attached debris.

### EdU Incorporation

Juveniles were incubated in a working concentration of 3 µM of EdU (Click-iT EdU Alexa Fluor 488 imaging kit, Invitrogen) in FSW for 1 hour, rocking at room temperature. The juveniles were then washed into 1:1 MgCl_2_: FSW for 10 minutes before being fixed in 4% PFA for 1 hour. Following fixation, the specimens were transferred to a glass three-well depression plate, washed into PBS, and incubated in PBS + 0.05% Triton-X-100 for 20 minutes, rocking at room temperature. They were then incubated with the commercial Click-it reaction mixture at room temperature for 30 minutes. The juveniles were washed into PBS and stored at 4°C until antibody labeling.

### Immunohistochemistry

Animals were incubated in a blocking solution of 10% heat-treated normal goat serum in PBS + 0.2% Triton-X-100 (PBT) for 45–60 minutes at room temperature. The anti-acetylated tubulin antibody (goat anti-mouse, Sigma-Aldrich Co.) was diluted 1:400 in blocking solution and incubated at 4°C overnight. Following incubation, the animals were washed four times, for 30 minutes each, in 0.2% PBT. A secondary antibody (goat anti-mouse, Invitrogen) was diluted 1:400 in block solution and added to animals for an overnight incubation at 4°C. Animals were once again washed four times, 30 minutes each, in 0.2% PBT. During the final two washes, Hoechst was added to PBT at a dilution of 1:1000. After the washes, the animals were cleared in 80% glycerol:20% PBS overnight at 4°C prior to imaging.

### Hindgut and whole-body length measurements

Animals were imaged on a Zeiss Axio Imager M2 with a red (594) filter set to visualize the anti-acetylated tubulin labeled cilia. The distance tool of Zeiss imaging software was used to measure the length of the whole body and of the ciliated hindgut. A ratio of hindgut length: whole body length was generated using the respective measurements tool. The box and whisker plot of ratio distributions was produced using Statistics Kingdom (Statistics Kingdom, 2017a).

### Replicates and Statistical analysis

When possible, uncut controls were sourced from the same broods as the cut experimental group (with exception of the 5 dpa larvae). At least 2 replicates were performed for each timepoint/treatment.

Statistical comparisons of ratios of hindgut length to whole-body length were made using the Shapiro-Wilk test (Statistics Kingdom, 2017c) was conducted on the various data sets to assign whether they fit a normal or non-normal distribution. It was shown that most of the data was not normally distributed. To compare the hindgut: whole body ratio of cut and uncut animals at the time points listed in Table 1, we conducted a Mann Whitney U (Statistics Kingdom, 2017b) statistical analysis.

### Microscopy and imaging

A Zeiss 710 confocal microscope, with 4x bidirectional and unidirectional scan settings for green and red lasers, was used to image the fluorescence of specimens. The resulting files were compiled into Z-stacks using ImageJ software (Schindelin et al., 2012). A Zeiss Axioskop II motplus compound microscope coupled with a SPOT FLEX digital camera (Diagnostic Instruments Inc.) was used to capture differential interference contrast (DIC) microscopy images. As noted in figure legends, multiple focal planes of DIC images were merged using Helicon Focus Software for some images (Helicon Focus 7 & 8). Images were cropped and adjusted (as a whole) for brightness and contrast in either Photoshop (Adobe Photoshop 2020) or Illustrator (Adobe Illustrator 2020). Figures were composed using the Adobe Illustrator software (Adobe Illustrator 2020).

### Whole mount in situ hybridization

Animals were collected either immediately following metamorphosis or at 7 days post metamorphosis, immobilized by incubation in a 1:1 MgCl: FSW solution for 10 minutes, then fixed in 4% paraformaldehyde in FSW at 4°C overnight. The animals were washed several times with PBS + 0.2% Triton and then slowly washed into 100% methanol and stored at -20°C. Whole mount in situ hybridization was conducted according to a previously published protocol (de Jong and Seaver, 2017). The DIG-labeled riboprobe was generated using the MegaScript T7 Transcription kit (Invitrogen) according to manufacturer instructions and stored in hybe buffer at -20°C. The *vasa* probe was used at a working concentration of 2 ng/µl and hybridized with animals at 65°C for 48 hours. The animals were incubated with an anti-digoxigenin-alkaline phosphatase conjugated antibody (Roche), such that the RNA probe can be visualized upon exposure to a color reaction solution with NBT/BCIP (nitro blue tetrazolium chloride/5-bromo-4-chloro-3-indolyphosphate) (Promega). The animals were visually monitored during development and all treatment groups were stopped concurrently, controls. Color development was stopped with extensive washes of PBT + 0.2% Triton and a final fixation using 4% PFA in FSW for 30 minutes. Tissues were cleared in 80% glycerol in 1x PBS before being mounted on microscope slides for analysis and imaging.

### Determination of segment number

The number of segments in a juvenile was determined by independently scoring three markers: ganglia in the nuclear stain, paired peripheral nerves in antibody staining, or chaetae. If two out of three markers scored, i.e., ganglia in the nuclear stain, paired peripheral nerves in antibody staining, or chaetae, yielded different results, then the common number was ruled the final number of total segments (frequently chaetae were fewer in number compared to presence of ganglia and peripheral nerves as chaetal development occurs very late in segment formation). On the rare occasion when all three markers yielded different segment numbers (e.g., 14,15,16), the following procedure was followed: each channel was scrutinized again and recounted. If still in disagreement, then an average of the three numbers was assigned as the final number of segments for that animal.

Changes in the appearance of the ventral nerve cord (vnc) were used to determine the boundary between pre-existing and regenerated segments. The ventral nerve cord was severed during amputation, such that there is a substantial decrease in thickness of the nerves as it crosses the amputation plane. Tissue distal to the position of this transition of the vnc were identified as newly regenerated.

## Acknowledgements

The authors thank Danielle de Jong for cloning the *C. teleta vasa* gene and generating the RNA probe used in this paper. The authors thank Brent Foster for critical reading of the manuscript.

